# The Genome In A Bottle HG002 assembly-based variant benchmark set enables comprehensive benchmarking of small and structural variants

**DOI:** 10.64898/2026.09.23.752440

**Authors:** Nathan D. Olson, Nathan Dwarshuis, Nancy F. Hansen, Sergey Koren, Arang Rhie, Jennifer H. McDaniel, Justin Wagner, Gwenn Berry, Kishwar Shafin, Adam English, Christopher T. Saunders, Ritu Kundu, James M. Holt, Donald Freed, Adetola A. Abdulkadir, Camille Daniels, Sina Majidian, Luis F. Paulin, Fritz J. Sedlazeck, Mark Fleharty, Yueyao Gao, Luca Denti, Lucas Brambrink, Andrew Carroll, Pi-Chuan Chang, Daniel E. Cook, Alexey Kolesnikov, Severine Catreux, Christopher Edlund, James Han, Lisa Murray, Yutong Qiu, Massimiliano Rossi, Konrad Scheffler, Sean Truong, Yina Wang, Rayan Chikhi, Carlos Flores, David Jáspez, José M. Lorenzo-Salazar, Adrián Muñoz-Barrera, Luis A. Rubio-Rodríguez, Mao-Jan Lin, Kavya Vaddadi, Ben Langmead, Chunlin Xiao, Sean K. McKenzie, Jonathan R. Belyeu, Michael A. Eberle, Zev Kronenberg, Juniper A. Lake, William J. Rowell, Aaron M. Wenger, Sayed M. Sahraeian, Omer Barad, Hila Benjamin, Ilya Soifer, Fereydoun Hormozdiari, Mohammed F. Mootor, Yash Patel, Takafumi N. Yamaguchi, Paola Bonizzoni, Paul C. Boutros, Adam M. Phillippy, Justin M. Zook

## Abstract

High-accuracy, high-resolution benchmark data are fundamental to improving the accuracy and generalizability of sequencing technologies and associated algorithms. Most existing benchmarks for DNA sequencing rely on mapping to a single linear reference, omitting inaccessible and highly repetitive complex genomic regions. Here, we present the Genome in a Bottle (GIAB) Consortium’s HG002 v5.0q benchmark set, a more comprehensive variant benchmark derived from a fully phased, highly curated, and accurate diploid assembly of HG002. Our approach leverages recent advances in long-read sequencing and assembly to provide confident small- and structural-variant calls across previously inaccessible regions, including segmental duplications, homopolymers, tandem repeats, and complex immune loci. The v5.0 benchmark significantly expands the callable genome relative to v4.2.1, adding ≈ 200 Mbp of sequence and increasing variant detection by 18 % for small variants and ≈3-fold for structural variants compared to previous benchmarks. The benchmark regions include more variants in repetitive regions, including 1.2× segmental duplications, 1.8× tandem repeats, and 2.0× homopolymers, compared to v4.2.1. Separate and combined small- and structural-variant benchmarks are available for three versions of the human reference genome (GRCh37, GRCh38, and CHM13). We validate the reliability of this benchmark set by comparing multiple sequencing technologies and variant calling pipelines and curating differences. The v5.0q benchmark provides a critical resource for the development and validation of next-generation sequencing technologies and analysis methods.

## Introduction

Aligning reads and calling variants against a reference genome is the most common method to identify biologically and clinically relevant variation. Benchmarks for germline small and structural variants have facilitated the development of sequencing technologies and bioinformatics tools,^1–4^ the training and testing of AI-based variant callers,^5,6^ and the validation of clinical tests.^7^ Previous whole-genome variant benchmarks have integrated multiple callsets from different technologies by either arbitrating between differing calls based on evidence of bias, such as those from the Genome in a Bottle Consortium (GIAB), or using pedigree-informed expectations of variant inheritance, such as Platinum Genomes.^8^ While these complementary approaches are effective for many regions of the genome, they can exclude or introduce errors around structural variants (SVs) and complex variants, where callsets may have differing variant representations.

Assembly-first analysis methods, in which a *de novo* assembly is generated from the reads rather than aligning them to a genome, enable the identification of variation that is particularly challenging or inaccessible with read-mapping-based methods due to mapping ambiguities and variants that are poorly represented in the VCF format. High-accuracy diploid T2T assemblies are enabling a new type of benchmark, a genome benchmark. In genome benchmarking, assemblies or inferred genomes (where a phased assembly is inferred from a set of variants and a reference genome) are compared directly to the benchmark genome assembly. Genome benchmarking is now possible with the T2T highly polished HG002v1.1 assembly and the new GQC tool from the T2T “Q100” project.^9^ With recent advances in sequencing technologies and assembly algorithms, essentially complete diploid human genomes can be generated using a combination of accurate long reads, ultralong reads, and phasing from trios, Hi-C, or Strand-Seq.^10,11^ While these diploid assemblies typically contain some errors in satellite repeats, variable number of tandem repeats (VNTRs), dinucleotide tandem repeats (TRs), and homopolymers, almost all errors were eliminated in the HG002v1.1 assembly.

As read-mapping-based variant calling remains the dominant analysis method in genomics, high-accuracy and comprehensive variant benchmarks are needed to advance the adoption of whole-genome sequencing in clinical settings and advance genomic research. To address these limitations, some specialized assembly-based variant benchmarks have been released.^12–14^ Assembly-based variant benchmarks focus on generating an accurate diploid genome assembly and then aligning it to the reference to produce phased variant calls, regardless of variant size. Building on previous work using synthetic diploid assemblies,^15^ GIAB has progressively released targeted assembly-based benchmarks for the MHC (major histocompatibility complex),^16^ challenging medically relevant genes,^12^ and chromosomes X and Y for HG002.^14^ Here, we use the highly polished HG002v1.1 assembly to create a comprehensive whole-genome variant benchmark, enabling robust benchmarking of variant calls.

To create this benchmark, we developed the `DeFrABB` (Development Framework for Assembly-Based Benchmarks) pipeline (https://github.com/usnistgov/defrabb), which takes any accurate diploid assembly as an input and outputs benchmark variants and regions on any reference, including GRCh37, GRCh38, and T2T-CHM13v2.0. This pipeline outputs phased SNVs, indels, and large insertions and deletions up to tens of Kbp, including many segmental duplications and TRs, challenging gene conversions, and complex variants. Benchmark sets generated using `DeFrABB` expand on GIAB’s previous specialized benchmarks for targeted regions and variant types by enabling benchmarking across the autosomes and sex chromosomes for small variants and SVs jointly or separately. To enable robust benchmarking, it excludes very large and complex SVs and copy number variants (CNVs), as these are regions for which no standards exist for representation or comparison. Using `DeFrABB` and the new HG002-T2TQ1000v1.1 assembly, we generated new GIAB HG002 v5.0q small- and structural-variant benchmark sets for GRCh37, GRCh38, and CHM13. We evaluated this benchmark with Miqa (https://www.magnalabs.co/), an online curation platform to curate putative errors from a variety of short- and long-read callsets, ensuring it reliably identifies errors.^17^ Because draft benchmarks were made available for public use and feedback, many publications have already demonstrated the utility of this benchmark for benchmarking challenging variants.^4,18–25^ Furthermore, as highly accurate assemblies become increasingly common, this pipeline could enable robust variant benchmarks for many genomes from different ancestries.

## Results

### Enabling reproducible, assembly-based benchmark generation

We created `DeFrABB` as a modular, open-source software tool to facilitate the generation and evaluation of draft variant benchmarks from high-quality diploid assemblies for the GIAB project. `DeFrABB` has three main components: assembly-based variant call generation, draft benchmark set generation, and evaluation (Fig. 1). The assembly-based variant benchmark also enables joint benchmarking of small variants, TRs, and SVs, which is important for a robust understanding of accuracy in regions with these variants.^18,20^ In addition, the assembly can be aligned to different references. Thus, we created comparable small- and structural-variant benchmark sets for GRCh37, GRCh38, and T2T-CHM13v2.0, enabling users to assess performance across different reference genome versions.

**Figure 1:**
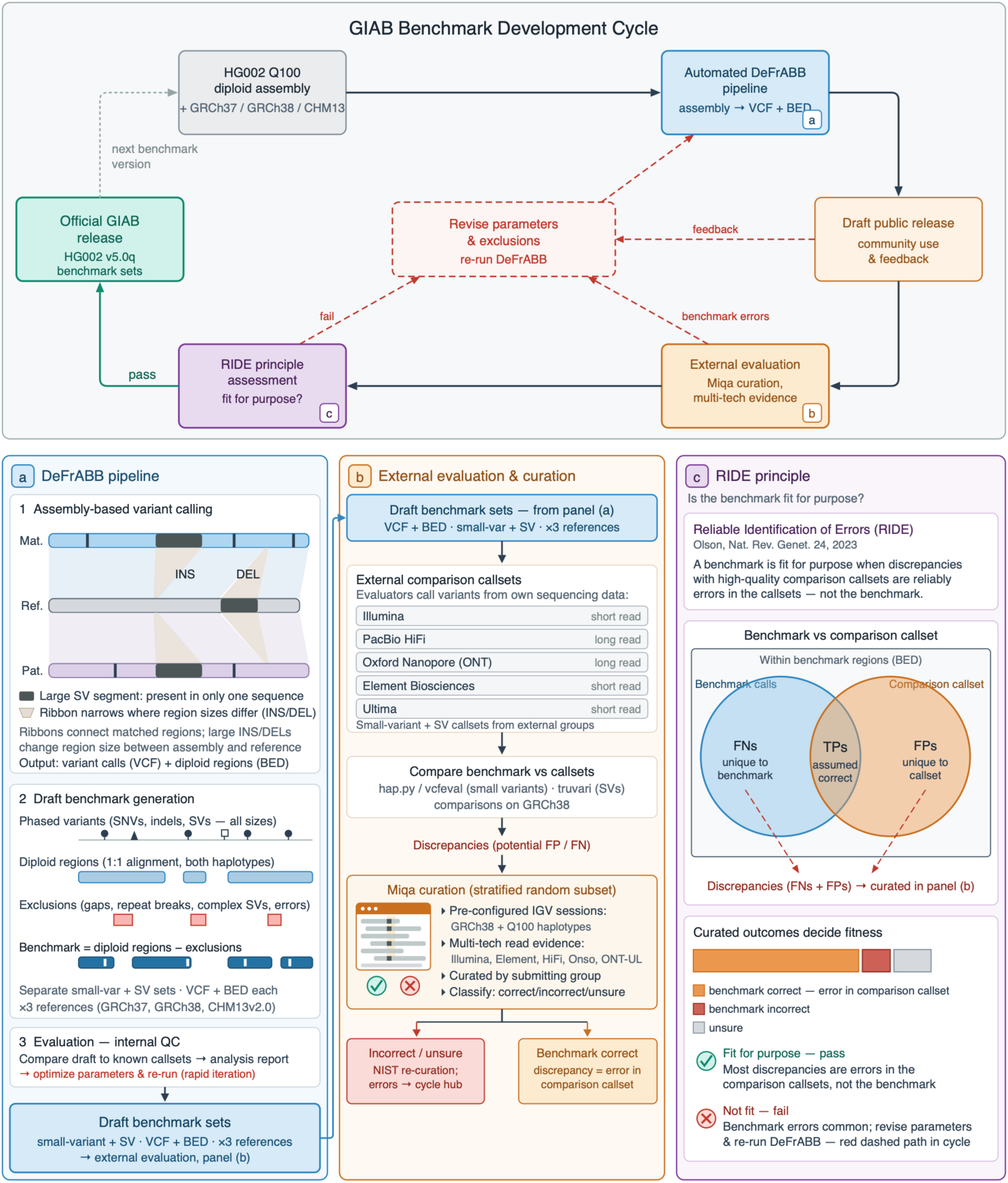
Development of the GIAB HG002 v5.0q whole-genome variant benchmarks. Top: the GIAB benchmark development cycle. The HG002 Q100 diploid assembly and reference genome (GRCh37, GRCh38, or CHM13) are inputs to the automated `DeFrABB` pipeline (panel a), which produces draft small variant and SV benchmark sets. Drafts are publicly released for community use, formally evaluated by external groups (panel b), and assessed against the RIDE principle (panel c). Red dashed paths mark checkpoints where community feedback, benchmark errors, or a failed assessment led to revised parameters and exclusions and a `DeFrABB` re-run; benchmarks that pass become official GIAB releases. (a) The three `DeFrABB` components. (1) Assembly-based variant calling: each haplotype is aligned to the reference; ribbons connect matched regions and narrow where large insertions (INS) or deletions (DEL) change region size, yielding variant calls (VCF) and diploid regions with 1:1 alignment of both haplotypes (BED). (2) Draft benchmark generation: benchmark regions are diploid regions minus exclusions (assembly gaps, alignment breaks in large repeats, complex SVs, and known assembly and variant-calling errors); phased variants of all sizes and benchmark regions are produced for small variants and SVs on all three references. (3) Evaluation: automated comparisons to existing callsets generate an analysis report used as internal QC for parameter optimization and rapid iteration. (b) External evaluation. Draft benchmarks (GRCh38) are compared with callsets contributed by external groups from multiple sequencing technologies using hap.py/vcfeval (small variants) and truvari (SVs). A stratified random subset of discrepancies is manually curated by each submitting group in the Miqa platform (Magna Labs, https://www.magnalabs.co) using pre-configured IGV sessions showing multi-technology read evidence on GRCh38 and both Q100 haplotypes; each discrepancy is classified as correct, incorrect, or unsure in the benchmark. Incorrect and unsure calls are re-curated at NIST and confirmed benchmark errors feed back into the development cycle. (c) The RIDE (“reliable identification of errors”) principle. A benchmark is fit for purpose when discrepancies with high-quality comparison callsets are reliably errors in the callsets rather than in the benchmark. Within benchmark regions (BED), calls unique to the benchmark are false negatives (FNs) and calls unique to a comparison callset are false positives (FPs); curated discrepancy outcomes determine whether the benchmark passes to official release.

### Benchmark content and genome coverage

The new v5.0q benchmark set covers more of the genome and more variants than previous benchmark sets, including previously inaccessible regions and more complex variants. The small variant benchmark set includes ≈ 700 000 more variants in GRCh38 than in v4.2.1, half of which are single-nucleotide variants (SNVs) and half are insertions/deletions (indels). The v5.0q benchmark set includes 1.5 % more of the autosomes than v4.2.1, as well as both sex chromosomes that were previously excluded. The v5.0q SV benchmark includes more than twice as many SVs on each of the 3 references as our previous benchmark, GIAB v0.6, which was only on GRCh37 (Table 1). The v5.0 SV benchmark set includes 226.1 % more variants in difficult genomic context (Fig. 2) and covers 9.1 % more of the reference genome. Coverage was higher for v5.0q than the previous benchmark for all chromosomes, except chromosomes 1, 8, and 16 for small variants (vs. v4.2.1, GRCh38) and chromosome 8 for SVs (vs. v0.6, GRCh37) (Fig. 3A). The lower coverage for chromosome 8 is due to exclusion of a common large (1.2 Mbp) inversion on the p arm of the chromosome (Fig. 3B). We took a conservative approach and excluded entire inversions from the benchmark because we were unable to define breakpoints confidently. A recent analysis found that most newly included variants in v5.0q were in repetitive regions.^26^ Qin and Li demonstrated that including these more challenging SVs is important for assessing SV caller accuracy, because most false positives (FPs) and false negatives (FNs) occur in these newly included regions. Even though only 1.2 % of the genome is in pangenome-derived low-complexity regions (excluding satellites), these repetitive regions contain 73.6 % of v5.0q SVs, and SVs in these regions have been shown to alter gene function and expression in some individuals with disease.

**Figure 2:**
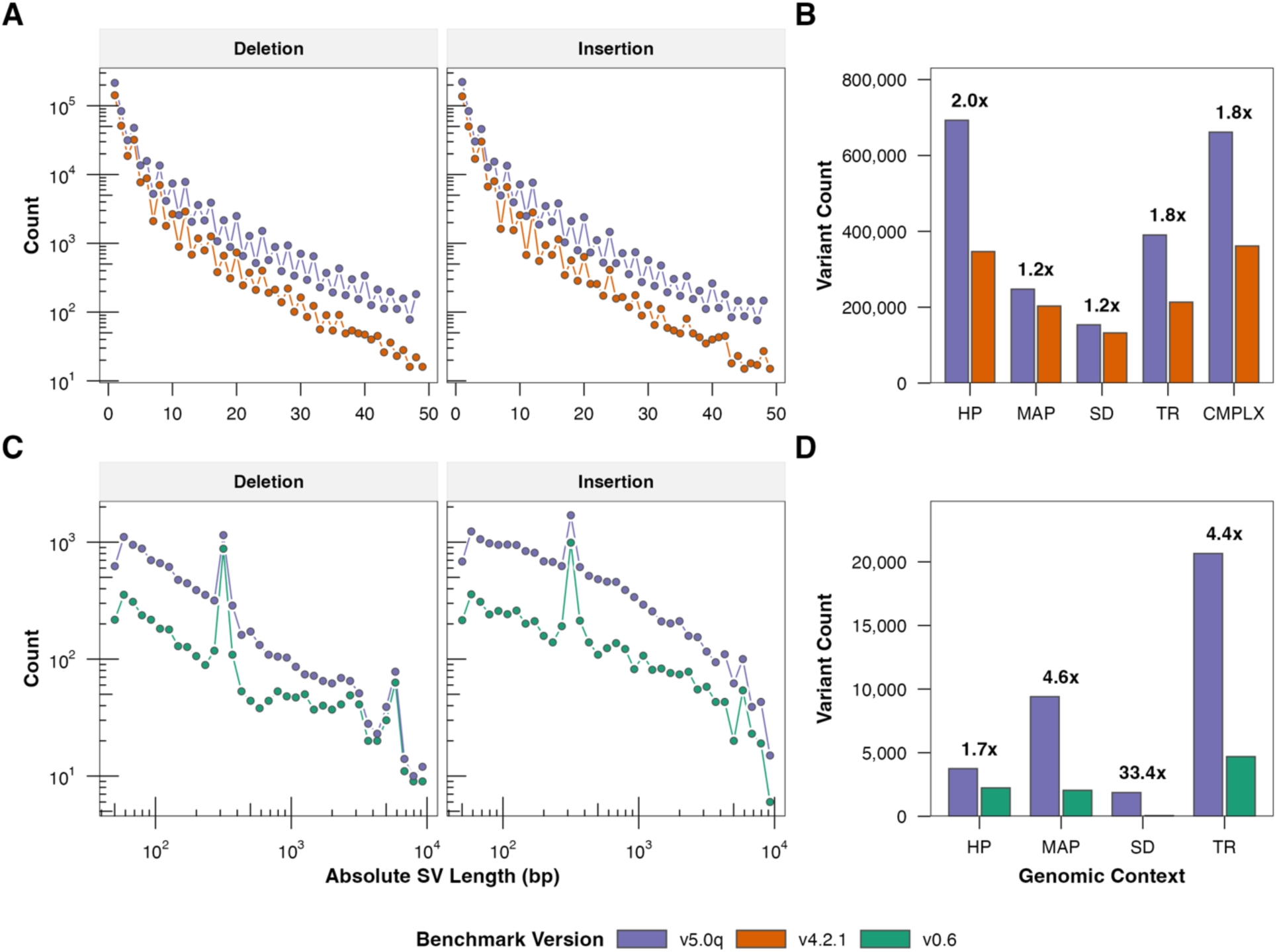
Comparison of the variant size distribution and number of variants in difficult genomic context between v5.0q and previous small variant (v4.2.1) and SV (v0.6) benchmark sets. Indels less than 50 bp (A) and SVs greater than 49 bp (C) for the small and SV benchmark sets, respectively. (B) The GRCh38 v5.0q small benchmark set variants compared to the v4.2.1 HG002 benchmark set variants. (D) The GRCh37 v5.0q structural benchmark set variants compared to the v0.6 HG002 SV benchmark set. HP: homopolymer; MAP: low-mappability; SD: segmental duplication; TR: tandem repeat.

**Figure 3:**
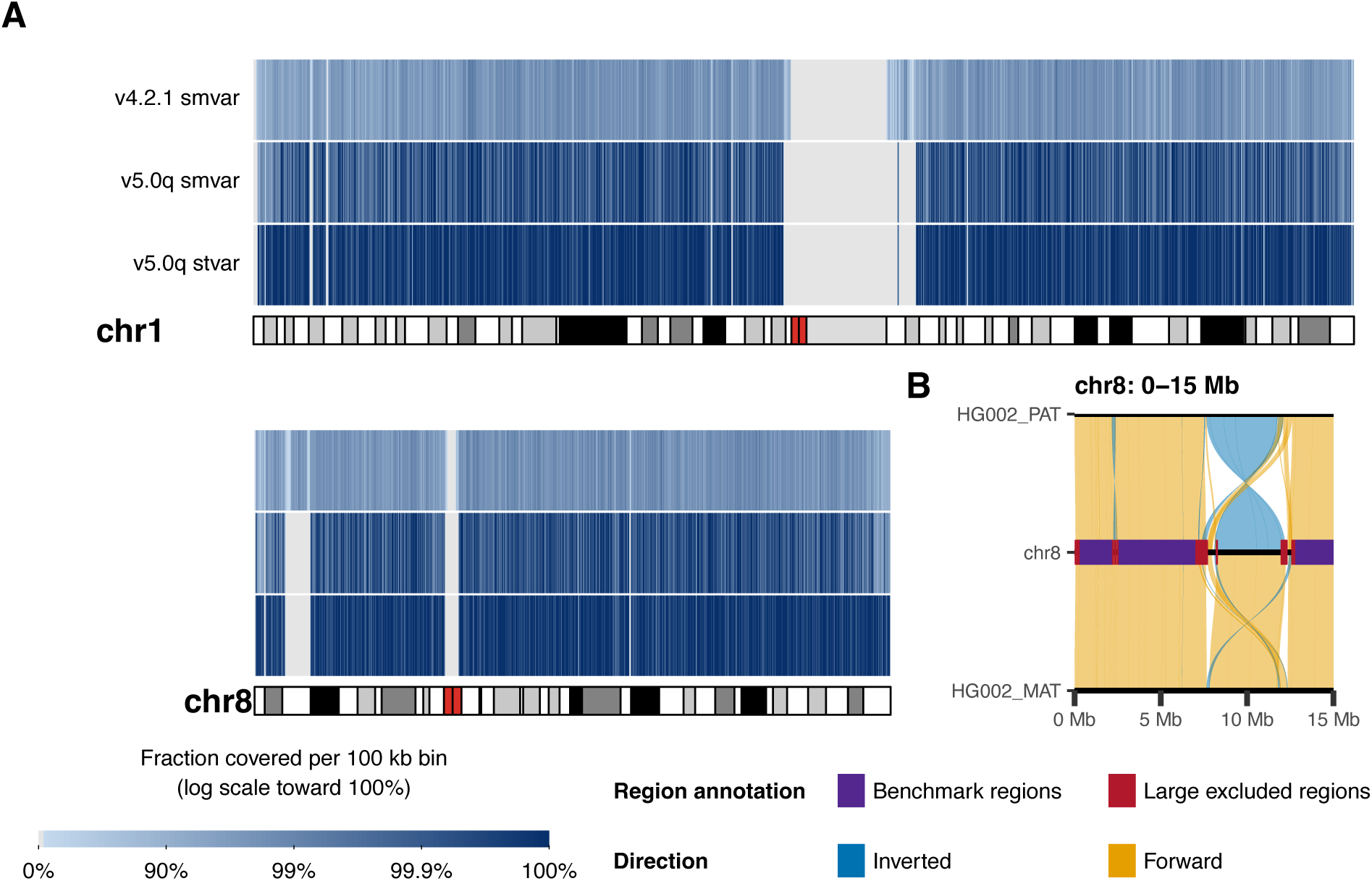
Benchmark region coverage of v5.0q benchmark region coverage relative to the v4.2.1 small-variant benchmark, for chromosomes 1 and 8 against GRCh38 (A). An excluded region around 10Mbp in chromosome 8 in v5.0q benchmark that was previously included in the v4.2.1 benchmark was excluded due to a large inversion between the reference assembly and the paternal haplotype (B). Coverage plots for all GRCh38 chromosomes are included as supplemental (Fig. S1).

**Table 1:** Comparison of v5.0q variant and regions statistics to previous GIAB HG002 small and structural variant benchmark sets. Summary of the v5.0q small variant and SV benchmark for HG002 on GRCh38. Small variant counts include single-nucleotide variants (SNVs) and insertions/deletions (INDELs) of less than 50 bp within benchmark regions. Altered bases is the number of SNVs plus the sum of the absolute length of the INDELs. Region (Mbp) is the total benchmark region size in megabases. % Ref - percentage of the assembled reference genome covered by benchmark regions. For the SV benchmark, variant counts include deletions (DEL) and insertions (INS) of 50 bp or larger within benchmark regions, and Variation is the sum of the variant lengths. Δ compares v5.0q (GRCh38) vs v4.2.1 (GRCh38) for small variants; v5.0q (GRCh37) vs v0.6 (GRCh37) for SVs. pp = percentage points.

| Type | Metric | V5.0Q | | | $\Delta$ vs previous version | |
| --- | --- | --- | --- | --- | --- | --- |
|  |  | GRCh37 | GRCh38 | CHM13v2.0 | n | % |
| Small Variant | SNV | 3 651 855 | 3 671 436 | 3 395 941 | +303 973 | +9.0 % |
|  | INDEL (< 50bp) | 951 308 | 954 482 | 885 454 | +395 476 | +70.7 % |
|  | Total Variants | 4 603 163 | 4 625 918 | 4 281 395 | +699 449 | +17.8 % |
|  | Altered Bases (Mbp) | 7.3 | 7.3 | 6.8 | +2.3 | +45.3 % |
|  | Region (Mbp) | 2 726 | 2 739 | 2 768 | +197 | +7.7 % |
|  | % of Reference | 95.4 % | 93.7 % | 88.8 % |  | +6.7 pp |
| SV | DEL (≥ 50 bp) | 10 561 | 10 852 | 12 841 | +6 491 | +159.5 % |
| | INS (≥ 50bp) | 17 571 | 17 763 | 12 204 | +12 024 | +216.8 % |
|  | Total SVs | 28 132 | 28 615 | 25 045 | +18 515 | +192.5 % |
|  | Variation (Mbp) | 19.1 | 18.6 | 16.1 | +11.8 | +161.3 % |
|  | Region (Mbp) | 2 742 | 2 756 | 2 820 | +229 | +9.1 % |
|  | % of Reference | 95.9 % | 94.3 % | 90.5 % |  | +8.0 pp |

In contrast to v0.6, which merged similar SV alleles, v5.0q provides exact phased sequence information, allowing comparisons at higher stringency than the defaults for tools like `truvari`.^27^ This enables assessment of base-level accuracy of SV calls, which can be important for certain TRs in genes. Despite the larger number of bases included on CHM13v2.0, benchmark set variant counts are smaller in CHM13v2.0 because (a) it fixes errors, and (b) it is of European genetic ancestry and therefore matches the genetic ancestry of HG002 more closely than the GRCh37 and GRCh38 references.

This benchmark captures additional challenging regions beyond the complex and interlocus gene-conversion-like loci described in our previous targeted benchmarks (e.g., Chin et al^28^). The *NEB* gene associated with myopathies has an 8.2 Kbp highly identical segmental duplication in the middle of the gene, with dense, small-scale variation on one HG002 haplotype, making it particularly challenging for short-read assessments. *POTEI* and *POTEJ*, part of a polymorphic gene family highly expressed in some germ cells and cancers,^29^ appear to undergo an interlocus gene-conversion-like event that causes mismapping of short and HiFi reads to *POTEJ* from *POTEI*. The *CLPS* gene is duplicated in a 12 Kbp insertion on one HG002 haplotype, and the *TAS2R43*/*PRH1* genes have many small variants and SVs.

*MUC3A* has a 7 Kbp to 8 Kbp TR expansion on each haplotype in the coding region, and there is a 58 Kbp to 63 Kbp insertion near the start of the adjacent *MUC12* gene, which is in our SV benchmark. As a particularly complex example, the *NPIP* gene family region on chromosome 16 exhibits complex SVs between individuals.^30^ While HG002 does not have inversions relative to GRCh38 in the *NPIPB6* to *NPIPB9* region, each haplotype harbors large insertions and/or deletions affecting genes. Both haplotypes have two additional tandem copies of *BOLA2B*+*SLX1A*+*SULT1A3* relative to GRCh38’s two copies, like CHM13 and most individuals, which is called as a ≈ 102 Kbp insertion in v5.0q SV benchmark (Fig. 4A). The paternal haplotype has a 13 Kbp insertion that duplicates *SULT1A1*, and the maternal haplotype has a 114 Kbp deletion of *NPIPB6* and the first copy of *EIF3C*. Both haplotypes have a large common inversion relative to CHM13, so that region is excluded from the CHM13-based benchmark.

**Figure 4:**
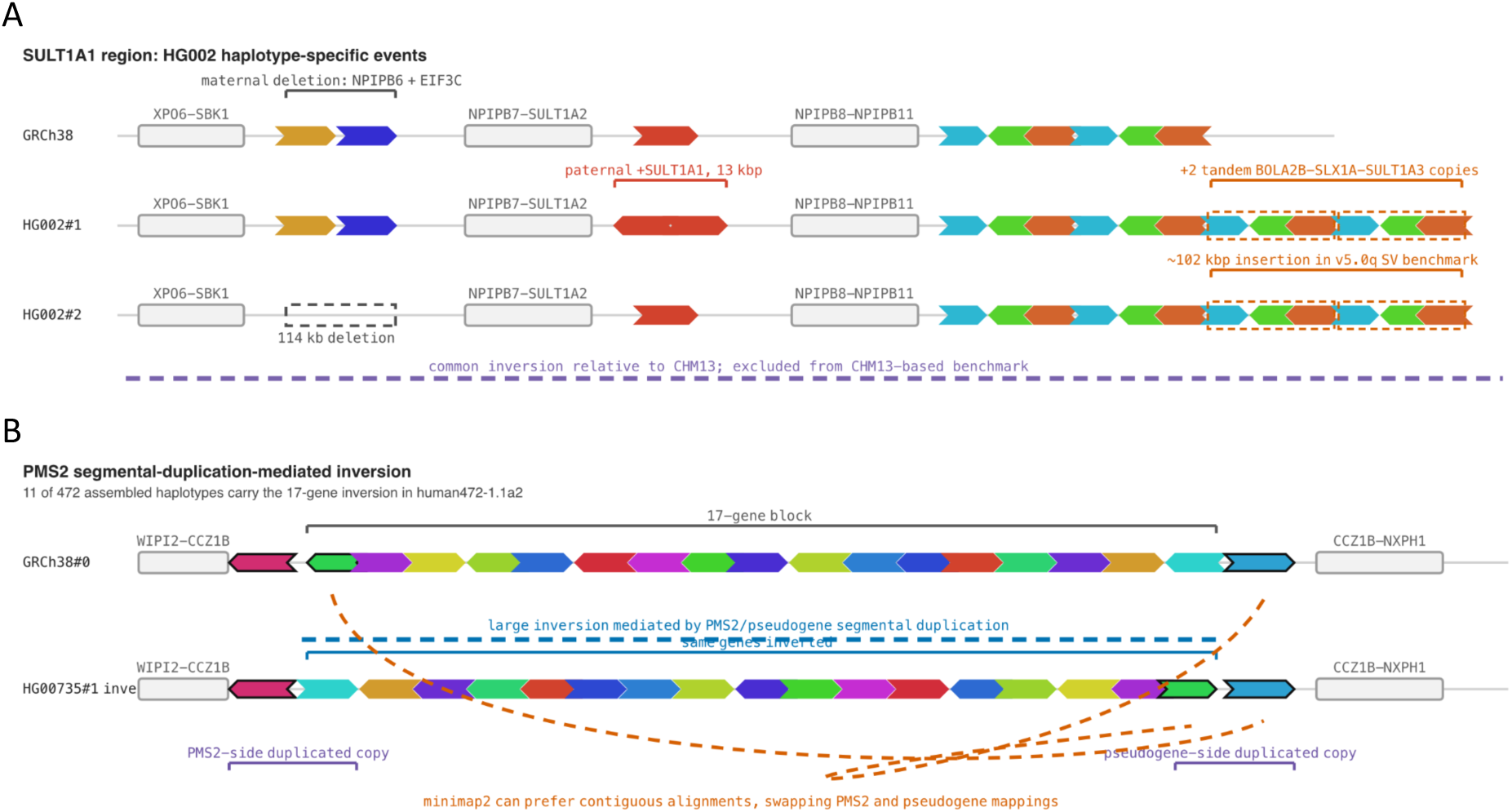
Pangene views of complex structural loci in HG002. (A) *SULT1A1* region haplotype-specific structural events, including a paternal 13 Kbp insertion duplicating *SULT1A1*, two additional tandem *BOLA2B–SLX1A– SULT1A3* copies (called as a ≈ 102 Kbp insertion in the v5.0q SV benchmark), and a maternal 114 Kbp deletion of *NPIPB6* and the first copy of *EIF3C*; a common inversion relative to CHM13 is also shown and is excluded from the CHM13-based benchmark. (B) *PMS2* segmental-duplication-mediated inversion, present in 11 of 472 assembled haplotypes in `pangene` human472-1.1a2, illustrates how the 17-gene inversion causes minimap2 to swap *PMS2*/pseudogene alignments. Diagrams are modified based on the *SULTA1* (https://pangene.bioinweb.org/view?graph=human472-1.1a2&gene=SULT1A1&step=3&ori=) and *PMS2* (https://pangene.bioinweb.org/view?graph=human472-1.1a2&gene=PMS2&step=10&ori=) `pangene` graphs.

Even with this nearly perfect HG002 assembly, we still exclude certain complex regions from the benchmark where no standards exist for variant representation or comparison. As with our previous targeted assembly-based benchmarks, we exclude *SMN1* and *SMN2* because they lie within a large repetitive region with complex SVs,^12^ though the variant calls in the genic regions appear reliable in our benchmark. Similar to our sex chromosome benchmark, we exclude the *DAZ* locus on chromosome X and the *TSPY2* locus on chromosome Y, which exhibit complex structural and copy-number variation.^14,31^ We investigated the *PMS2* segmental duplication in this benchmark and found a large inversion on one haplotype, mediated by the segmental duplication containing *PMS2* and its pseudogene. This inversion is in 11 of 472 assembled haplotypes in `pangene` human472-1.1a2 (Fig. 4B) and results in inversion of 17 genes.^32^ Because minimap2 prefers more contiguous alignments, it maps the inverted *PMS2* to its pseudogene and vice versa, resulting in many small and large variants in these regions. This was excluded from our benchmark due to an inversion and a break in alignment within the segmental duplication, highlighting the challenges in representing variants in this region. Unlike assembly alignments and ultralong reads, *PMS2* reads < 30 Kbp are generally mapped to *PMS2* rather than the pseudogene and miss the inversion.

### External Evaluation and Validation

The benchmark sets were compared to variant callsets contributed by external groups to validate that the benchmark sets are fit for purpose and adhere to RIDE, 5 small variant callsets and 7 SV callsets (Table 2). A random subset of discrepancies was selected from different complex strata for each callset. The external evaluators manually curated 40 variants for each submitted SV callset and 70 per small variant callset. Variants classified as incorrect or unsure in the benchmark set were re-curated at NIST; the external evaluators confirmed final curation classifications (Fig. 5). To verify that the benchmark sets can reliably identify errors, a binomial generalized linear model (GLM) was used to estimate the probability that a discrepancy between the benchmark and a comparison callset is an error in the comparison callset (i.e., the benchmark is correct). The outcome was binarized from the curation: yes = correct in the benchmark; no/unsure = not confirmed correct. Separate models were fit for small and SVs. As the goal was to establish a lower bound on benchmark reliability, one-sided 95 % lower confidence bounds are reported. The lower confidence bound of the estimates for all variant types and contexts was greater than 82.7 % for the small variant benchmark set and 68.1 % for the SV benchmark set.

**Figure 5:**
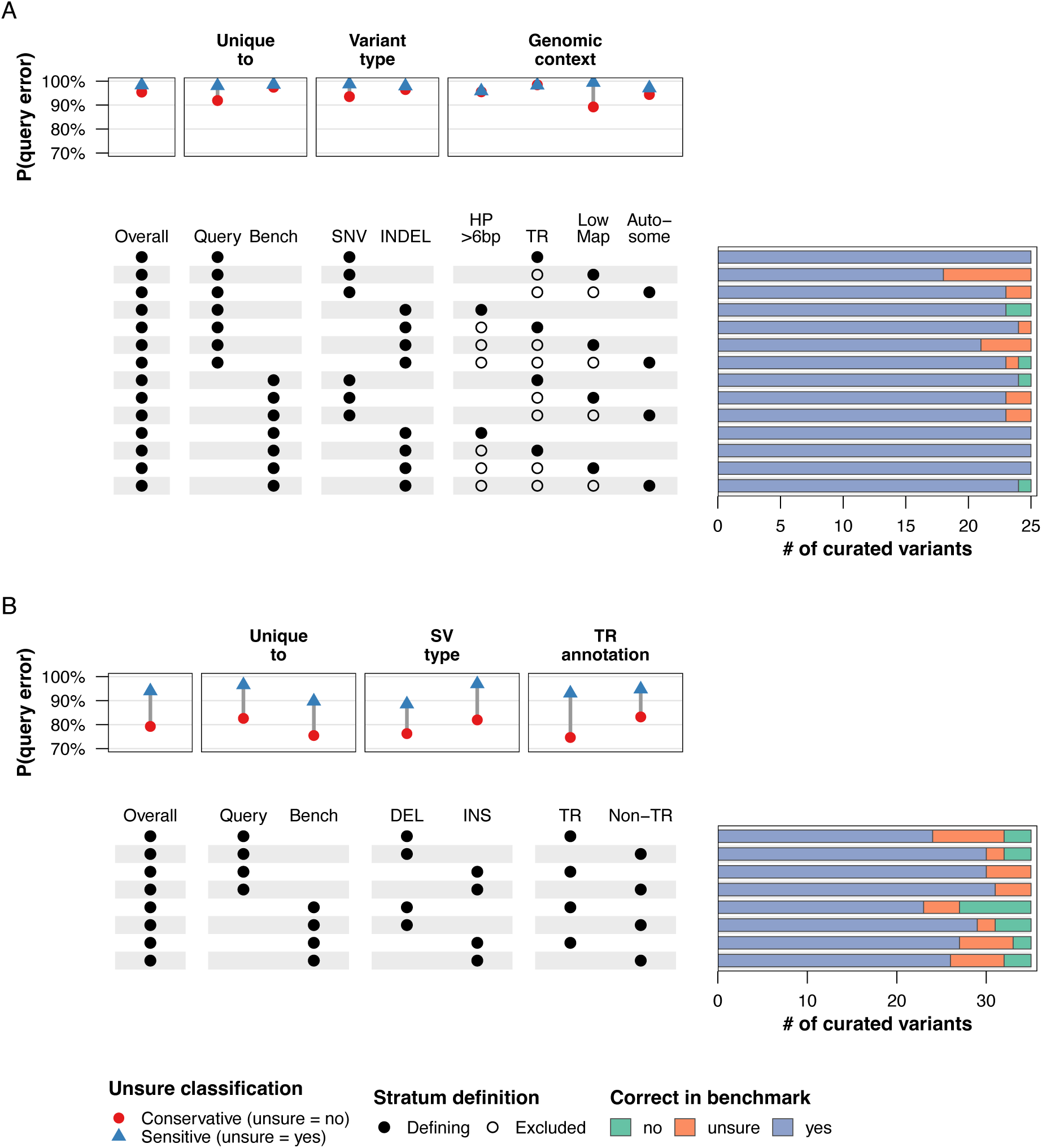
External evaluation summary. Number of manually curated variants as correct (yes, no, or unsure) and estimated probability that a discrepancy between the Q100 v5.0q HG002 benchmark and a comparison callset represents an error in the comparison callset, stratified by variant class and genomic context for the (A) small variant benchmark sets by strata and (B) SVs. Each row represents the feature categories defining the strata sampled for variant curation, along with the curations. The points in the plots above the feature categories are marginal probability estimates from bias-reduced logistic regression models with proportional weighting. Red circles indicate the conservative assumption (unsure-classification variants treated as callset errors); blue triangles indicate the sensitive assumption (unsure variants treated as callset correct). The grey segment spanning each row shows the range of estimates under the two assumptions, reflecting the uncertainty introduced by classifying unsure variants as correct or incorrect. The Overall estimate is not conditioned on any stratum.

**Table 2:**
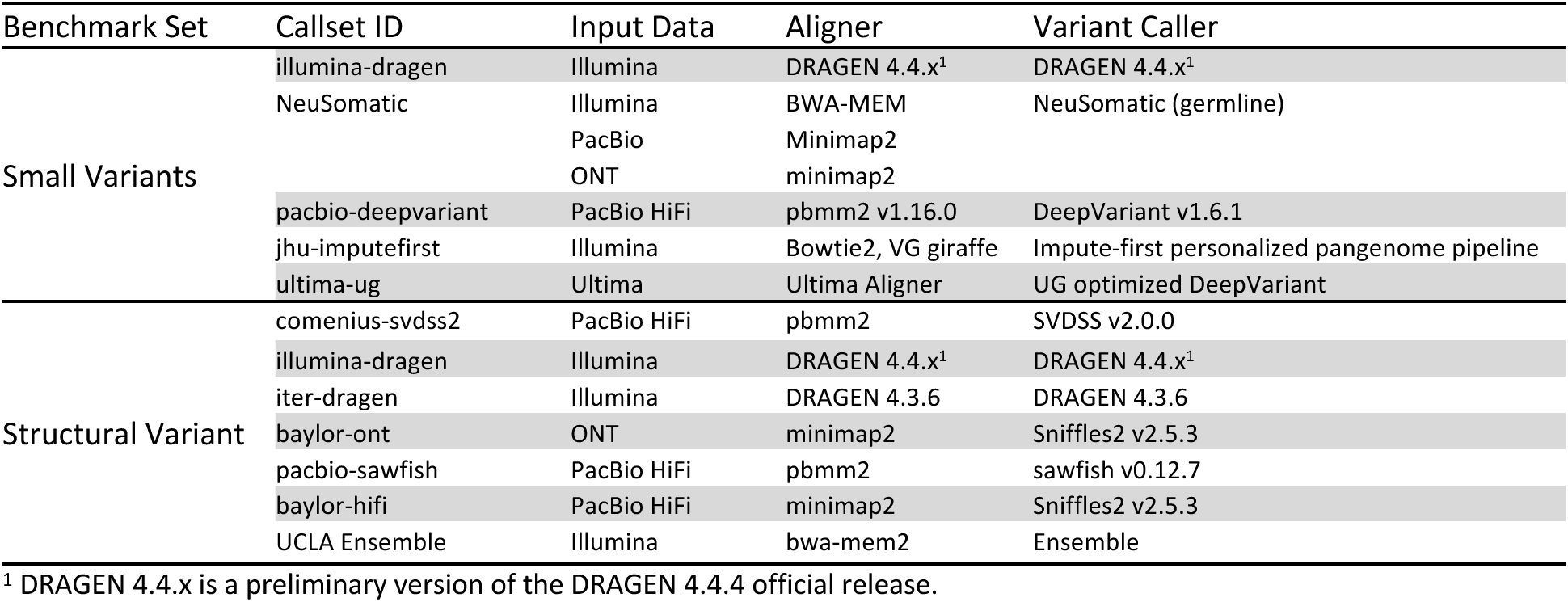
External Callsets Used for v5.0q Benchmark Evaluation. Small variant and structural variant callsets contributed by external groups for comparison and evaluation of the v5.0q benchmark sets on GRCh38.

| Benchmark Set | Callset ID | Input Data | Aligner | Variant Caller |
| --- | --- | --- | --- | --- |
| Small Variants | illumina-dragen | Illumina | DRAGEN 4.4.x<sup>1</sup> | DRAGEN 4.4.x<sup>1</sup> |
|  | NeuSomatic | Illumina | BWA-MEM | NeuSomatic (germline) |
|  |  | PacBio | Minimap2 |  |
|  |  | ONT | minimap2 |  |
|  | pacbio-deepvariant | PacBio HiFi | pbmm2 v1.16.0 | DeepVariant v1.6.1 |
|  | jhu-impute-first | Illumina | Bowtie2, VG giraffe | Impute-first personalized pangenome pipeline |
|  | ultima-ug | Ultima | Ultima Aligner | UG optimized DeepVariant |
| Structural Variant | comenius-svdss2 | PacBio HiFi | pbmm2 | SVDSS v2.0.0 |
|  | illumina-dragen | Illumina | DRAGEN 4.4.x<sup>1</sup> | DRAGEN 4.4.x<sup>1</sup> |
|  | iter-dragen | Illumina | DRAGEN 4.3.6 | DRAGEN 4.3.6 |
|  | baylor-ont | ONT | minimap2 | Sniffles2 v2.5.3 |
|  | pacbio-sawfish | PacBio HiFi | pbmm2 | sawfish v0.12.7 |
|  | baylor-hifi | PacBio HiFi | minimap2 | Sniffles2 v2.5.3 |
|  | UCLA Ensemble | Illumina | bwa-mem2 | Ensemble |

<sup>1</sup> DRAGEN 4.4.x is a preliminary version of the DRAGEN 4.4.4 official release.

### Comparison to Platinum Pedigree Benchmark

Our benchmark complements and improves upon the recent Platinum Pedigree (PP) benchmark, which substantially expanded upon the GIAB v4.2.1 benchmark for HG001 by using short and long reads from a large family pedigree.^33^ We found the PP and GIAB v5.0 benchmarks are similar in content for small variants, including ≈ 4.6 million (v5.0q) and ≈ 4.3 million (PP) small variants spanning ≈ 2.7 Gbp of GRCh38 (Fig. 6). Our assembly-based approach results in much less fragmented benchmark regions, with only 9 658 regions in the SV benchmark and 28 797 regions in the small variant benchmark, vs. over 1 million regions in the PP benchmark. This fragmentation of benchmark regions can cause problems when benchmarking the complex variants common in repetitive regions, which can have different representations.^34^

**Figure 6:**
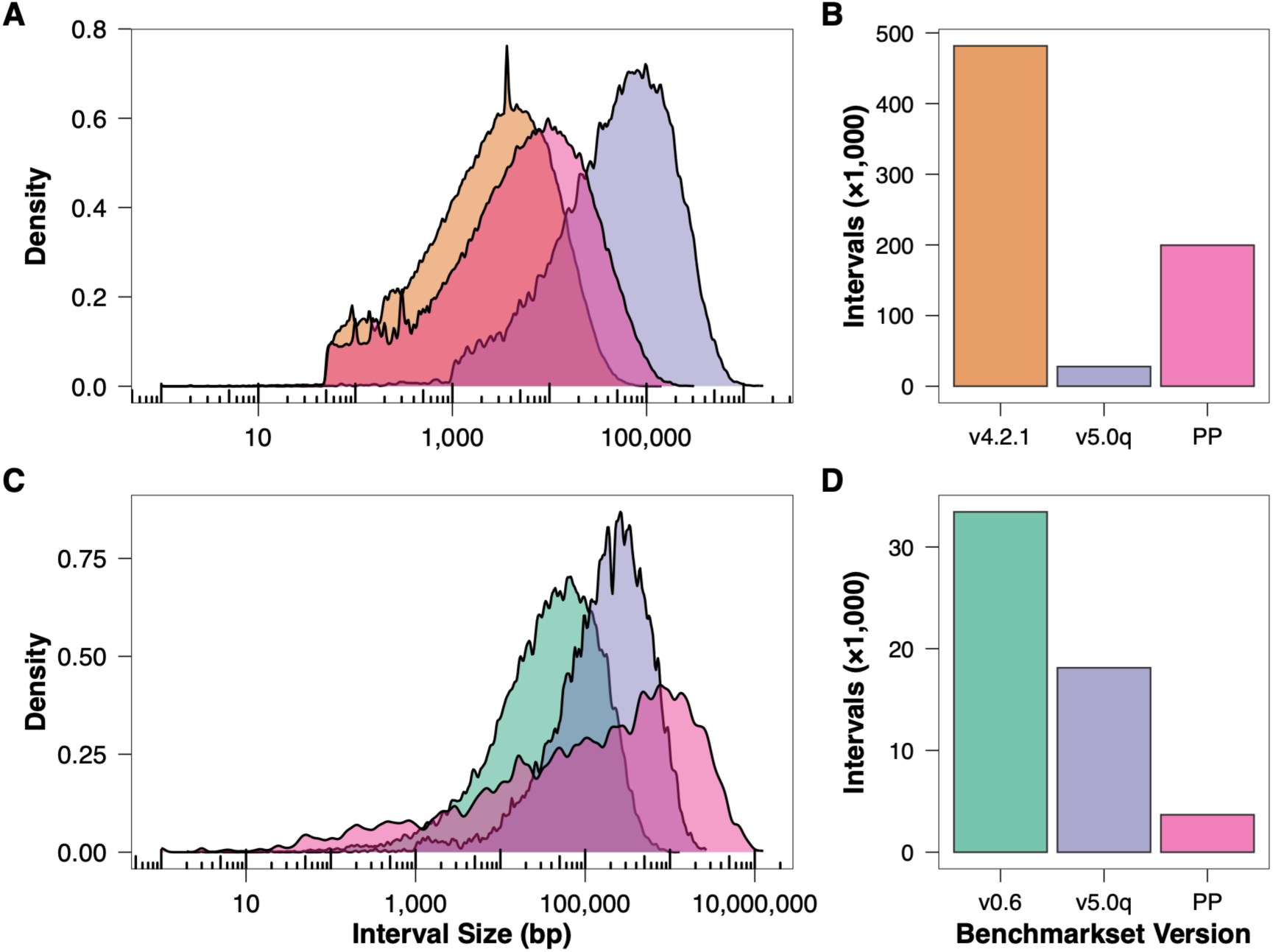
**Benchmark region interval comparison** for previous and v5.0q GIAB small and SV benchmark sets to Platinum Pedigree NA12878. Small variant benchmark region interval size distribution (A) and number of intervals (B). SV benchmark region size distribution (C) and counts (D).

While the blood DNA-based sequencing used for PP enables a more robust understanding of *de novo* variants without cell line artifacts, the NIST Reference Material and renewable cell line resource may differ from the blood in some places, causing challenges in benchmarking new technologies or a laboratory’s own sequencing process with the Platinum Pedigree calls. In our benchmark, we exclude likely *de novo* and mosaic variants in the HG002 cell line to minimize the cell line’s limitations (Table 3).^35^ We found that our assembly-based approach for identifying these excluded all recently identified mosaic variants in HG002, which can vary in frequency between cell line batches.^35^ We expect the new GIAB and PP benchmarks to be complementary for users, given their differences in benchmark construction, DNA source, donor genetic ancestry, sex, and filtering processes.

**Table 3:** Genome context excluded from v5.0q benchmark regions.

| Exclusion | Description | Reason |
| --- | --- | --- |
| Partially contained Segmental Duplications | Regions annotated as segmental duplications in reference assembly that are not fully contained in diploid aligned regions. | Assembly-based variant calls are inaccurate or ambiguously aligned due to large complex SVs. |
| Partially contained Tandem Repeats | Regions annotated as tandem repeats > 10 Kbp in reference assembly that are not fully contained in diploid aligned regions | Assembly-based variant calls are inaccurate or ambiguously aligned due to large complex SVs. |
| Partially contained Satellites | Regions annotated as satellites in reference genome that are not fully contained in diploid aligned regions. | Assembly-based variant calls are inaccurate or ambiguously aligned due to large complex SVs. |
| PAV-inversions | Inversions called by PAV. | Inversions are sometimes called as nearby insertions and deletions by dipcall, though variants inside large inversions may still be correct. |
| Flanks | 15 Kbp upstream and downstream of assembly alignment breakpoints. | Assembly-based variant calls are inaccurate or ambiguously aligned due to large complex SVs |
| VDJ | VDJ region. Somatically variable in the cell line. | Low confidence in the assembly. |
| HG002Q100v1.1 errors | Low confidence positions in HG002 Q100 v1.1 assembly. Identified during manual curation and polishing of the assembly. | Low confidence in the assembly. |
| TSPY2 | TSPY2 segmental duplication. | Assembly can be aligned in two alternate ways due to complex SVs. |
| Consecutive SVs | Intervals in the assembly-assembly alignment represented by an insertion immediately followed by a deletion or a deletion immediately followed by an insertion. Individual insertions and deletions are > 10 bp each. | Variants are missing due to errors in dipcall VCF. |
| Gaps | Gaps in reference assembly (Ns > 50 bp). 16S rDNA operons for CHM13. | Variant calls cannot be confidently called in these regions. |
| Dipcall-bugs T2TACE | Manual identified errors in dipcall due to a bug in dipcall. | Errors in dipcall VCF. |
| SV regions<sup>1</sup> | Structural variants (≥ 50 bp) and tandem repeats containing SVs (based on truvari anno trf TRFstart and TRFend annotations). Intervals within 5 bp merged. For intervals overlapping regions in all TR and homopolymer, overlapping intervals are also excluded. 50 bp of slop added to the resulting intervals then merged with a maximum distance of 1 Kbp. | Small variant benchmarking tools unable to reliably identify errors. |

<sup>1</sup> Only for small variant benchmark regions

## Discussion

Because early drafts of the v5.0q benchmarks were made public for community feedback, many publications have already demonstrated their utility. Critical feedback from a variety of use cases helps ensure the robustness of the benchmarks. The draft benchmarks were used in the development and validation of several new variant calling tools, including demonstrating that `sawfish` performs well for larger SVs and lower coverages,^19^ `DNAscope` using short and long reads has tens of thousands fewer small variant errors,^4^ `gcSV` and blend-seq using short and long reads reduce SV errors,^37,38^ `DRAGEN` has a 30 % increase in recall for SVs in v4.4 vs. v4.3,^39^ an impute-first approach to pangenomes can produce more accurate variants,^40^ and using a pangenome reduces errors by 5 % for small variants with DeepVariant.^21^ As described above, the SV benchmark was shown to enhance performance evaluation in challenging, repetitive regions substantially and to resolve them precisely. The combined phased small variants and SVs enabled the development of two new benchmarking tools, `vcfdist2` and `aardvark,` that jointly benchmark phased small variants and SVs, a capability shown to be important in many repetitive regions with complex variants.^18,41^ In addition to benchmarking, our benchmark has been used to train the `DeepVariant` 1.9.0 model to call more complex small variants.^42^ These efforts demonstrate the value of making these draft benchmarks openly available to the community as we worked towards the final versions.

As benchmarks become more comprehensive, stratification by genome context and variant type becomes increasingly important. For example, methods focused on analyzing coding regions may not need to be accurate in most very long homopolymers or satellite DNA. Alternatively, while the precise length of polyA tails in Alus and LINEs may not be important for many applications, the length of the much smaller number of G- or C-homopolymers in promoters might be important.

This work is a companion to a manuscript proposing the HG002 Q100 assembly itself as a ‘genome benchmark’.^9^ In the genome benchmarking paradigm, assembly evaluation is simpler and more comprehensive when evaluating assembly sequences directly rather than converting them to variant calls for comparison with variant benchmarks. Conversely, phased variant calls can be converted into a variant-constructed genome assembly and compared to the Q100 assembly as an alternative to using the variant benchmarks in this work. In regions assessed by the variant benchmark, the FP, FN, and genotype error metrics may be more relevant to most downstream variant-based applications than the current performance metrics from genome benchmarking. However, the genome benchmarking approach complements our variant benchmarks by enabling assessment of variant calls outside the variant benchmark regions. These regions were excluded due to large structural or copy-number differences between one or both HG002 haplotypes and the reference. Genome benchmarking can assess variants in these regions, though reconstructing genomes from them is challenging. More work is needed, using pangenomics or other specialized tools, to evaluate and compare complex differences among individuals in these regions. Future work might also enable inclusion of some of these regions in variant benchmarks if standards for variant representation and comparison were developed.

## Data and Code availability

- All benchmark VCFs, BEDs, and auxiliary files are hosted on the GIAB FTP under https://ftp.ncbi.nlm.nih.gov/ReferenceSamples/giab/release/AshkenazimTrio/HG002_NA24385_son/v5.0q/.
- HG002 Q100 v1.1 assemblies are deposited at NCBI (GCA_018852605.3, GCA_018852615.3) and mirrored at https://github.com/marbl/hg002.
- DeFrABB pipeline: https://github.com/usnistgov/defrabb
- Manuscript results, figures, and tables: https://github.com/nate-d-olson/q100-variant-benchmark-paper

## Acknowledgements

The authors would like to thank Drs. Patricia Kiesler, Craig Greenberg, and Myeshia Shelby for feedback on the draft manuscript. This work would not be possible without the incredible, collaborative efforts of the T2T and GIAB consortia. NDO would like to thank Bob Wier and Phil Lesh for the long, strange trip. PB has received funding from MSCA-RISE PANGAIA Horizon 2020 n. 872539. L.D. has received funding from the European Union’s Horizon program under the Horizon Europe grant agreement (ASVA-CGR No.∼101180581). S.M. is supported by the SciLifeLab & Wallenberg Data Driven Life Science Program (grant: KAW 2024.0159). M.L., K.V., and B.L.’s contributions were supported by the National Human Genome Research Institute grant R01HG011392 to B.L. P.C.B.’s contributions were funded by grants from the National Cancer Institute (R01CA270108, U2CCA271894, U24CA248265). F.J.S. receives research support from PacBio, Illumina, and Oxford Nanopore Technologies. This work was supported in part by the National Center for Biotechnology Information of the National Library of Medicine (NLM), National Institutes of Health (NIH). N.F.H., S.K., A.R., and A.M.P. were supported by the Intramural Research Program of the National Human Genome Research Institute, National Institutes of Health. C.X. was supported by the Intramural Research Program of the National Library of Medicine, National Institutes of Health. N.D.O., N.D., J.H.M., J.W., and J.M.Z. were supported by the National Institute of Standards and Technology.

K.S., L.B., A.C., P.C., D.C., and A.K. are employees of Google LLC and own Alphabet stock as part of the standard compensation package. S.C., C. E., J.H., L.M., Y.Q., M.R., K.S., S.T., and Y.W. are employees of Illumina and own stock as part of a standard compensation package. S.K.M. is an employee and shareholder of Oxford Nanopore Technologies. J.R.B., M.A.E., Z.K., J.A.L., W.J.R., and A.M.W. are employee’s and shareholders of PacBio.

The contributions of the NIH author(s) are considered Works of the United States Government. The findings and conclusions presented in this paper are those of the author(s) and do not necessarily reflect the views of the NIH, the U.S. Department of Health and Human Services, NIST, or the United States Government. Certain equipment, instruments, software, or materials are identified in this paper in order to specify the experimental procedure adequately. Such identification is not intended to imply recommendation or endorsement of any product or service by NIST, nor is it intended to imply that the materials or equipment identified are necessarily the best available for the purpose. The manuscript text and code were authored with the assistance of Claude models from Anthropic and GPT models from OpenAI. AI was used to refine language, improve clarity, and enhance readability in accordance with the authors’ instructions. Code was similarly drafted and debugged with AI. All content, scientific claims, and conclusions have been reviewed and verified by the authors to ensure accuracy and originality.

## Methods

### V5.0q Benchmark Set Generation

#### HG002 Assembly

GIAB and T2T selected HG002 for the aspirational Q100 project because the individual and his parents were broadly consented by the Personal Genome Project for its extensive public resources. Specifically, HG002 has high-coverage public genomic data from essentially all sequencing technologies. Lymphoblastoid (GM24385) and two induced pluripotent stem cell lines (GM26105 from B-lymphocyte and GM27730 from fibroblast) are publicly available in the Coriell Cell line repository. In addition, NIST has developed Reference Materials 8391 and 8392 from large homogeneous batches of DNA, and numerous commercial products for clinical validation use HG002 as the background DNA.

The Q100v1.1 assembly underwent several rounds of polishing and manual corrections to fix most errors. However, some ambiguous regions remained, including long homopolymers that could not be phased to one haplotype, some VNTRs, and highly repetitive ribosomal DNA regions. These issues were identified manually and by mapping reads to the assembly, and they were lifted to GRCh37, GRCh38, and CHM13 and excluded from this benchmark. We also conservatively continued to exclude some errors in v1.0 that were likely fixed in v1.1.

#### Development Environment for Assembly-Based Benchmarks (DeFrABB)

The `DeFrABB` pipeline has three main components: assembly-based variant calling, draft- benchmark generation, and draft-benchmark set evaluation (Fig. 1A). `DeFrABB` uses the Snakemake workflow language engine, which enables the development of transparent and reproducible data analysis pipelines that can be run across multiple environments. `DeFrABB` runs are configurable using the `resources.yaml` file, where users can define the input diploid assemblies, reference genomes, and paths to other input files, as well as which exclusions to apply when generating the draft benchmark bed. A separate `analysis.tsv` file is used to define the individual draft benchmark sets and evaluations. Pipeline dependencies are defined as conda environments.

For the assembly-based variant callset generation component, the initial set of variant calls and diploid regions is defined by comparing the two haplotypes of a diploid assembly to a reference genome assembly. To generate the variant calls, the individual haplotypes are aligned to the reference assembly using assembly-assembly alignment. The assembly-assembly alignments are then used to identify differences (variants) between the user-provided diploid assembly and the reference genome. The assembly-assembly alignments are also used to identify diploid regions, regions of the genome with a one-to-one alignment with the reference genome for the two diploid assembly haplotypes. The variant calls and diploid regions form the basis of the draft benchmark set’s variants and regions. The assembly-based variant caller and parameters are configurable. The `dipcall`^15^ and `PAV`^43^ assembly-based variant callers are currently supported. The `DeFrABB` modular structure allows for the inclusion of other variant callers or variant-type-specific callers in the future. The assembly-based variant calls are then processed for use as benchmark variants. VCF processing steps include variant formatting (e.g., normalization using `BCFtools`) and annotation (using `Truvari`).

The next main component of `DeFrABB` is draft benchmark generation, where the assembly-based variant calls and diploid regions are used to generate draft benchmark sets. Here, the VCFs are processed for use with variant-benchmarking tools, such as `hap.py` (for small variants) and `Truvari` (for SVs). For example, fixing genotypes in XY non-autosomal regions and normalizing HG002. The benchmark regions are defined by excluding the user-defined genomic context. Errors and biases in genome assembly and variant calling, reference assembly biases, VCF format limitations, and limitations of variant benchmarking tools can all lead to incorrect comparisons when benchmarking variant calls and are therefore excluded from the benchmark regions.^17^ To exclude variants with representation issues, not correctly handled by benchmarking tools, a self-discrepancy step is performed where the draft benchmark set is compared to itself. All FPs and FNs from the self-comparison are excluded.

`DeFrABB` has rules for excluding genomic context based on assembly-based variant calls, diploid-assembled regions, reference assembly annotations, variant characteristics (e.g., SVs in small variant benchmark sets), or user-provided bed files.

Evaluation is the final component of `DeFrABB`, in which the draft benchmark sets are compared with known high-quality variant callsets. `DeFrABB` uses the small-variant benchmarking tool `hap.py` to evaluate small-variant benchmark sets and `Truvari` for SV benchmark sets. Combined with summary statistics for the assembly-based variant calls and draft benchmark sets, the evaluation results are summarized in an analysis report. This analysis report serves as an initial QC step, allowing for parameter optimization and rapid iteration.

`DeFrABB` was used to create the v5.0q small variant and SV benchmark sets from the HG002-Q100v1.1 assembly for three different versions of the human reference genome assembly: GRCh37, GRCh38, and CHM13. The assembly-based variant caller `dipcall` was used to generate the initial variant calls and diploid regions. To increase mapping around SVs and the MHC region, the following z parameter was used: `-z200000,10000`. The resulting VCFs were annotated using the `truvari anno` modules `svinfo, trf, remap`, and `repmask`. Variants in the small variant benchmark set were normalized using `BCFtools` (1.14). `BCFtools` was also used to convert genotypes in non-pseudoautosomal regions of chrX and chrY.

The diploid assembled regions were refined to define the v5.0q benchmark regions. The refinement steps included the exclusion of genomic regions with either systematic assembly errors or assembly-based variant-calling errors, as well as regions of the genome that present challenges for variant representation and comparisons (Tables 3 and 4). We started with regions where there was a 1:1 alignment between each assembled haplotype and the reference (except for chromosomes X and Y).

**Table 4:** Bases removed from benchmark regions by exclusion for GRCh38. Number of bases in the excluded genomic context overlapping the diploid assembled regions and the percent of diploid assembled regions. Individual exclusion regions may overlap; the total number of bases excluded is less than the sum of individual exclusions.

| Exclusion | Bases Excluded | % of dip.bed |
| --- | --- | --- |
| Segmental Duplications | 52 311 253 | 1.841 |
| SV regions (small variant benchmark only) | 29 069 865 | 1.023 |
| PAV-inversions | 23 518 680 | 0.828 |
| Satellites | 16 627 547 | 0.585 |
| Flanks | 10 220 302 | 0.360 |
| Tandem Repeats | 6 553 662 | 0.231 |
| Gaps | 6 172 312 | 0.217 |
| HG002Q100v1.1 errors | 5 039 842 | 0.177 |
| VDJ | 3 105 977 | 0.109 |
| HG002-mosaic | 1 178 158 | 0.041 |
| TSPY2 | 589 097 | 0.021 |
| dipcall-pav_discrep-smvar | 333 608 | 0.012 |
| Consecutive SVs | 251 531 | 0.009 |
| dipcall-pav_discrep-stvar | 127 132 | 0.004 |
| Dipcall-bugs T2TACE | 45 604 | 0.002 |
| Self-Discrepancies (small variant benchmark only) | 15 177 | 0.001 |

Then we excluded gaps in the assembly and their flanking sequences (rDNA sequences for CHM13), as well as any large repeats > 10 Kbp (satellites, TRs, and segmental duplications) that had a break in the assembly-to-reference alignment on either haplotype. We also exclude regions affected by SVs from the small variant benchmark bed. Segmental duplications and satellites within 40 Kbp are merged before checking for alignment breaks in repeat regions, to conservatively exclude nearby satellites and segmental duplications if a break occurs within a satellite or segmental duplication. Additional exclusions were applied for known `dipcall` bugs, discrepancies between `PAV` and `dipcall` callsets, and HG002 assembly errors and mosaic variants. The HG002 assembly errors corrected in v1.1 of the Q100 and will likely be included in the next benchmark version. This polished, highly accurate assembly enables the inclusion of many challenging, repetitive regions that were previously excluded from benchmarks. However, some regions are still excluded from our benchmark due to reference issues, errors, and ambiguities in aligning the assembly to the reference, challenges in representing variation relative to a linear reference, and errors or mosaic variants in the assembly.

#### External Evaluation

The draft benchmark sets were evaluated against 5 small and 7 structural-variant callsets contributed by external groups (Table 2). Methods used to generate comparison callsets are listed in alphabetical order below.

##### Baylor HiFi and ONT

HG002 data PacBio HiFi data was downloaded from PacBio’s data portal (https://www.pacb.com/connect/datasets/) for the Application Note of the GIAB trio HG002-4 (Variant detection, assembly, epigenetics) https://downloads.pacbcloud.com/public/revio/2022Q4/. The three replicates were merged into a single file and down-sampled to 30× coverage using `SAMtools view` (seed=15485863).^49^ ONT PromethION R10 FlowCell with the 5khz mode Dorado basecaller (Date: 2023/07/06) sequenced at the Human Genome Sequencing Center, Baylor College of Medicine, with an average coverage of 80x. The same alignment and variant calling methods were used for both the PacBio HiFi and ONT data. Raw reads were aligned to the GRCh38.p13 reference genome using `minimap2` (2.28-r1209),^47^ and variant calling was performed with `Sniffles2` v2.5.3^50^ using the following commands.

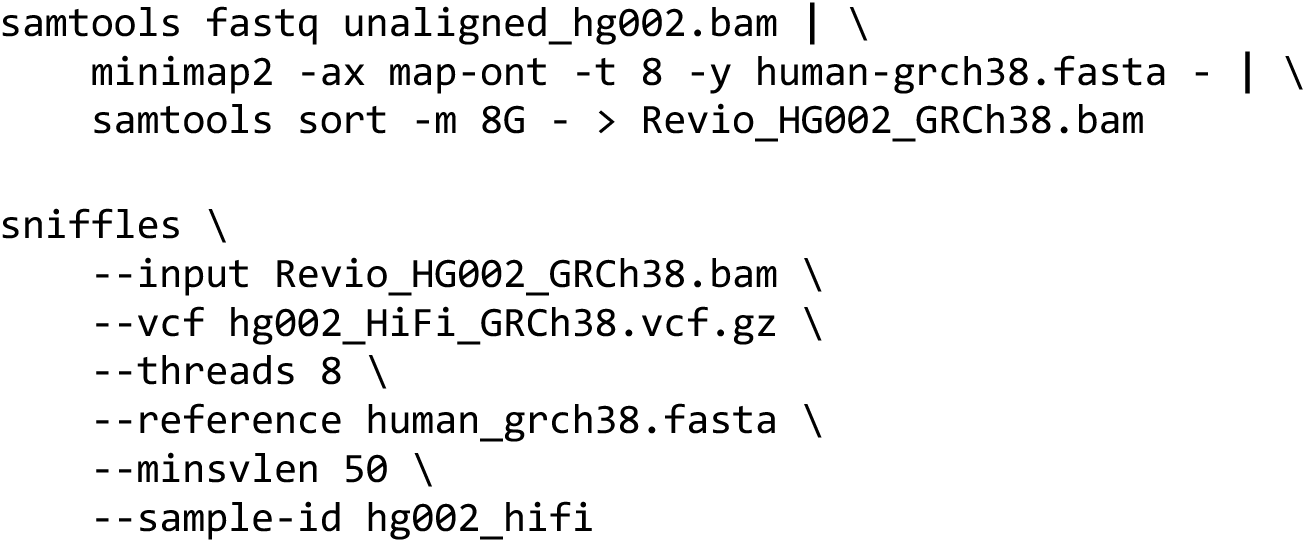

##### Comenius SVDSS2

`SVDSS` is an SV caller specially tailored to PacBio HiFi data.^51^ It uses an algorithm based on specific substrings to increase sensitivity compared to other callers and produce haplotype-resolved SVs. To curate the HG002 structural variant callset, we used version 2.0.0, which improves on previous v1.0.x versions by more clearly separating alternate alleles across the two haplotypes. `SVDSS2` indeed uses alignment-based haplotagging (based on short variants) to distinguish the two haplotypes better and precisely identify SV alleles, even at multiallelic non-reference loci. Since the SVDSS genotyping module is still undergoing validation, we considered all calls provided (even those with genotype 0/0). GRCh38 reference genome was downloaded from ftp://ftp.ncbi.nlm.nih.gov/genomes/all/GCA/000/001/405/GCA_000001405.15_GRCh38/seqs_for_alignment_pipelines.ucsc_ids/GCA_000001405.15_GRCh38_no_alt_analysis_set.fna.gz. PacBio HiFi sample of HG002 (SRA accession id: PRJNA586863) was downloaded from the Human Pangenome Reference Consortium (Sequel II System, Chemistry 2.0, size selection with SageELF to 20 Kbp, ≈ 16-fold coverage). Raw reads (runs m64011_190830_220126 and m64011_190901_095311) were directly downloaded from the HPRC repository (https://s3-us-west-2.amazonaws.com/human-pangenomics/working/HPRC_PLUS/HG002/raw_data/PacBio_HiFi/20kb/). The HG002 reads were aligned to the GRCh38 reference genome with pbmm2 (v1.16.0, installed from conda) using the CCS preset and successively merged into a single BAM file with SAMtools^49^using the following commands.

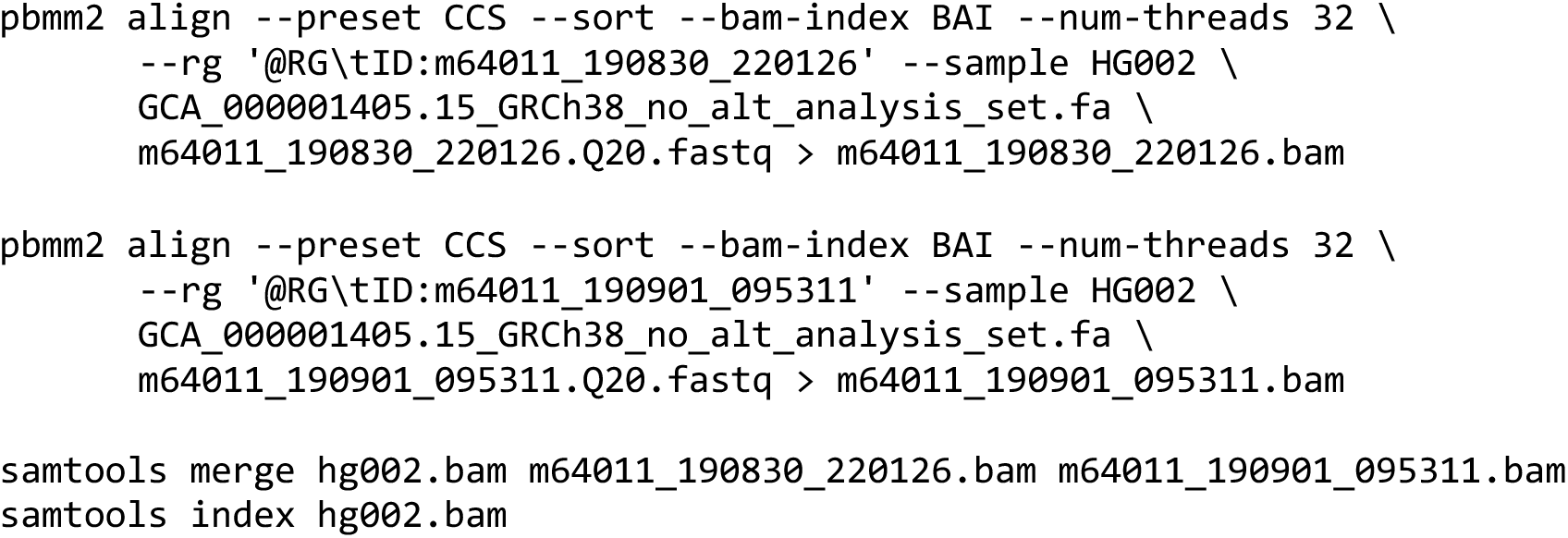

We first called short variants using `DeepVariant` (v1.6.1, ran via singularity)^6^ and then haplotagged the input alignments using `WhatsHap` (v2.3, installed from conda).^52^ Using the haplotagged alignments, we called structural variations using the `SVDSS`^51^ pipeline (https://github.com/Parsoa/SVDSS/releases/tag/v2.0.0). We considered all alignments with a minimum mapping quality of 20 (`--min-mapq parameter`) and an accuracy of 0.02 or less (`--acc parameter`, defined as the ratio of mismatches to matches in the alignment). We called insertions and deletions longer than 50 bp `(--min-sv-length parameter`) and supported by at least 2 reads (default value for `--min-cluster-weight` parameter).

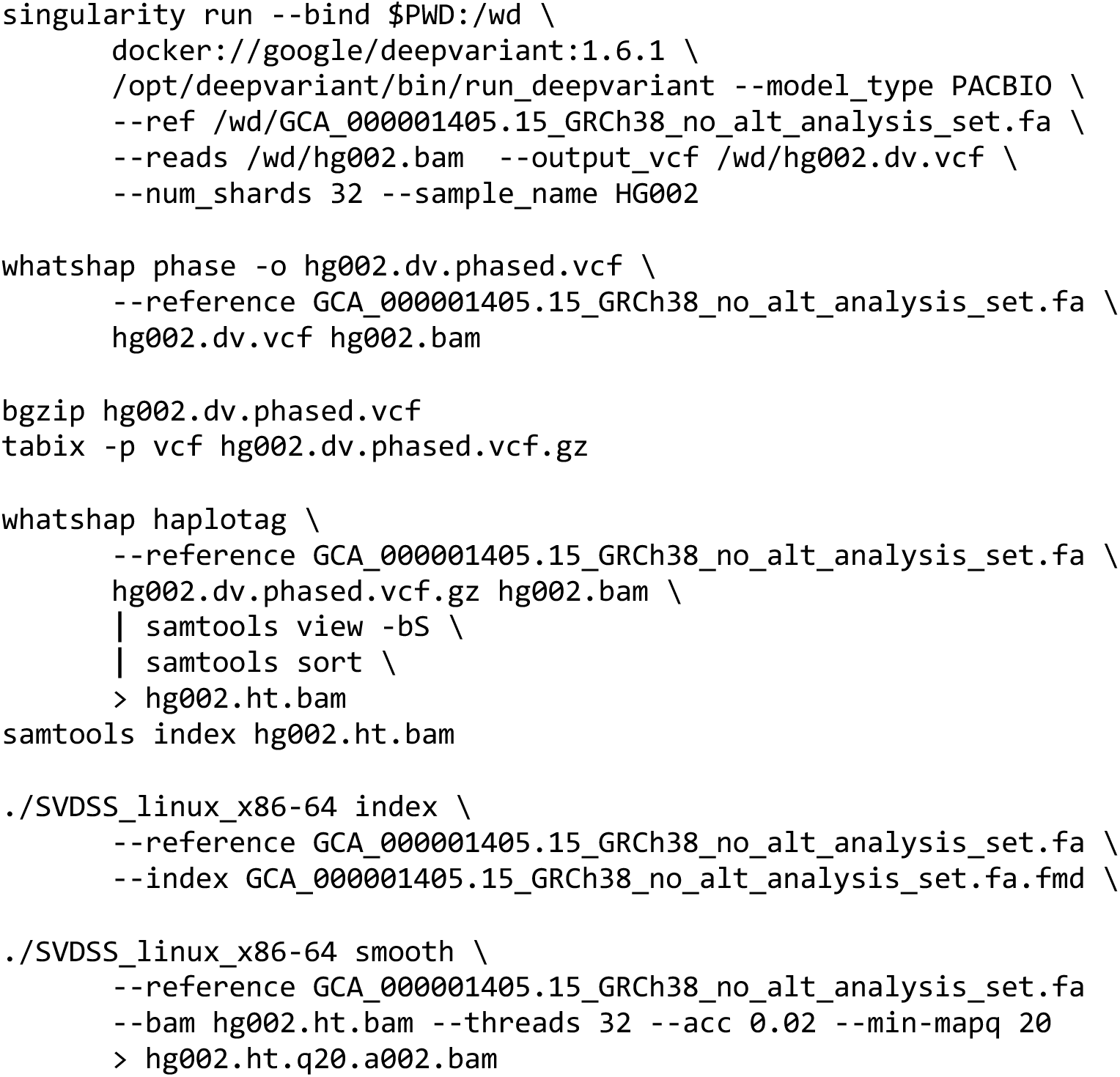
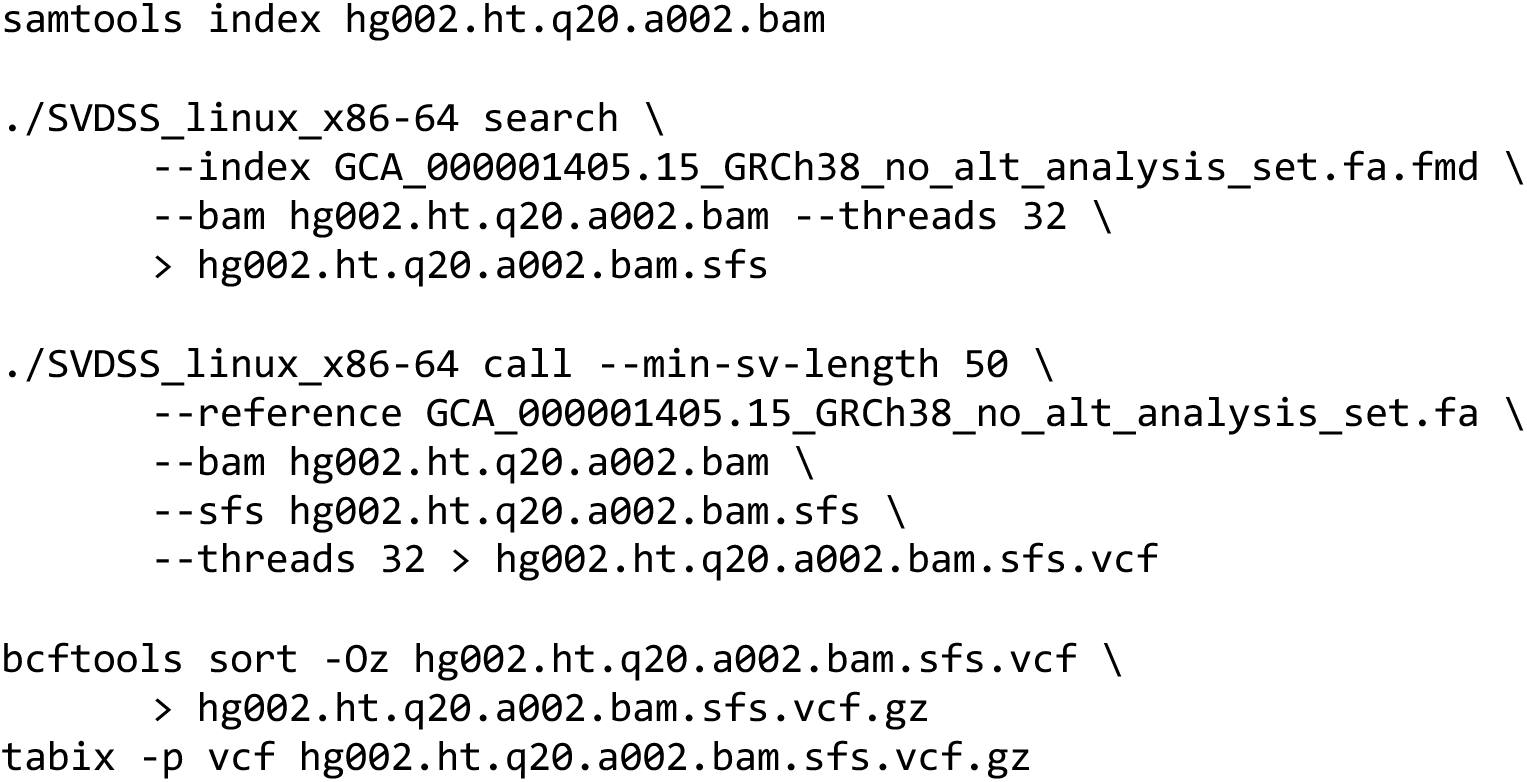

##### Illumina DRAGEN (Small and Structural Variants)

A NA24385 (HG002) sample sourced from the Coriell Institute For Medical Research was prepared using the TruSeq DNA PCR-Free library prep kit, and sequenced on the NextSeq 2000 P4 flow cell, using 2 × 151 bp read length, XLEAP-SBS reagents, and the HP Advanced Custom Recipe (https://www.illumina.com/science/genomics-research/articles/hp-advanced-custom-recipe.html). The HG002 paired-end FASTQ sample was aligned and mapped with a pre-release version of `DRAGEN` 4.4.x (available early 2025). ^3^ A single command-line call was used to align the paired-end FASTQs and call variants (https://help.dragen.illumina.com/dragen-v4.3/product-guide/dragen-v4.3/dragen-recipes/dna-germline-wgs.) `SNV_OPTION` and the `SV_OPTIONS` were configured to enable the detection of small and structural variants, respectively. The reference used for read alignment and mapping is the DRAGEN hg38 multigenome (graph) reference hash table (https://s3.us-east-1.amazonaws.com/webdata.illumina.com/downloads/software/dragen/references/genome-files/hg38-alt_masked.cnv.graph.hla.methyl_cg.rna-11-r5.0-1.tar.gz). For detailed methods about the individual variant callers and options, see https://help.dragen.illumina.com/dragen-v4.3/product-guide/dragen-v4.3/dragen-dna-pipeline/small-variant-calling and https://help.dragen.illumina.com/dragen-v4.3/product-guide/dragen-v4.3/dragen-dna-pipeline/sv-calling

##### ITER DRAGEN

The input data consists of paired-end short-read FASTQ files from the HG002 reference sample, publicly available through the National Institute of Standards and Technology (NIST) Genome in a Bottle (GIAB) project ^53^ (https://opendata.nist.gov/pdrsrv/mds2-2336/input_fastqs/HG002.novaseq.pcr-free.35x.R1.fastq.gz and https://opendata.nist.gov/pdrsrv/mds2-2336/input_fastqs/HG002.novaseq.pcr-free.35x.R2.fastq.gz). The sequencing depth is approximately 35×, and the data were generated using PCR-free library preparation on an Illumina NovaSeq platform. Read alignment and mapping were performed using the `DRAGEN`^3^ pipeline version 4.3.6 in a single integrated step. The reference genome used was the `DRAGEN` hg38 graph-based reference, publicly available from Illumina as https://webdata.illumina.com/downloads/software/dragen/resource-files/hg38-alt_masked.cnv.graph.hla.rna-10-r4.0-1.tar.gz. This pangenome-based reference improves the mapping accuracy of Illumina reads in “Difficult-to-Map Regions” of the genome and enhances downstream variant calling by extending the reference genome with alternative variant paths derived from a sample cohort. During the mapping step, duplicate reads were marked with the `--enable-duplicate-marking parameter`, and the aligned reads were written to a sorted BAM file.

Structural variant (SV) calling was performed simultaneously with read alignment using the `DRAGEN` pipeline version 4.3.6. While the analysis was primarily focused on SV detection, it also included configurations for small-variant calling, copy-number variants (CNVs), HLA typing, targeted calling, and Variable Number Tandem Repeats (VNTRs). The analysis was performed using the following command.

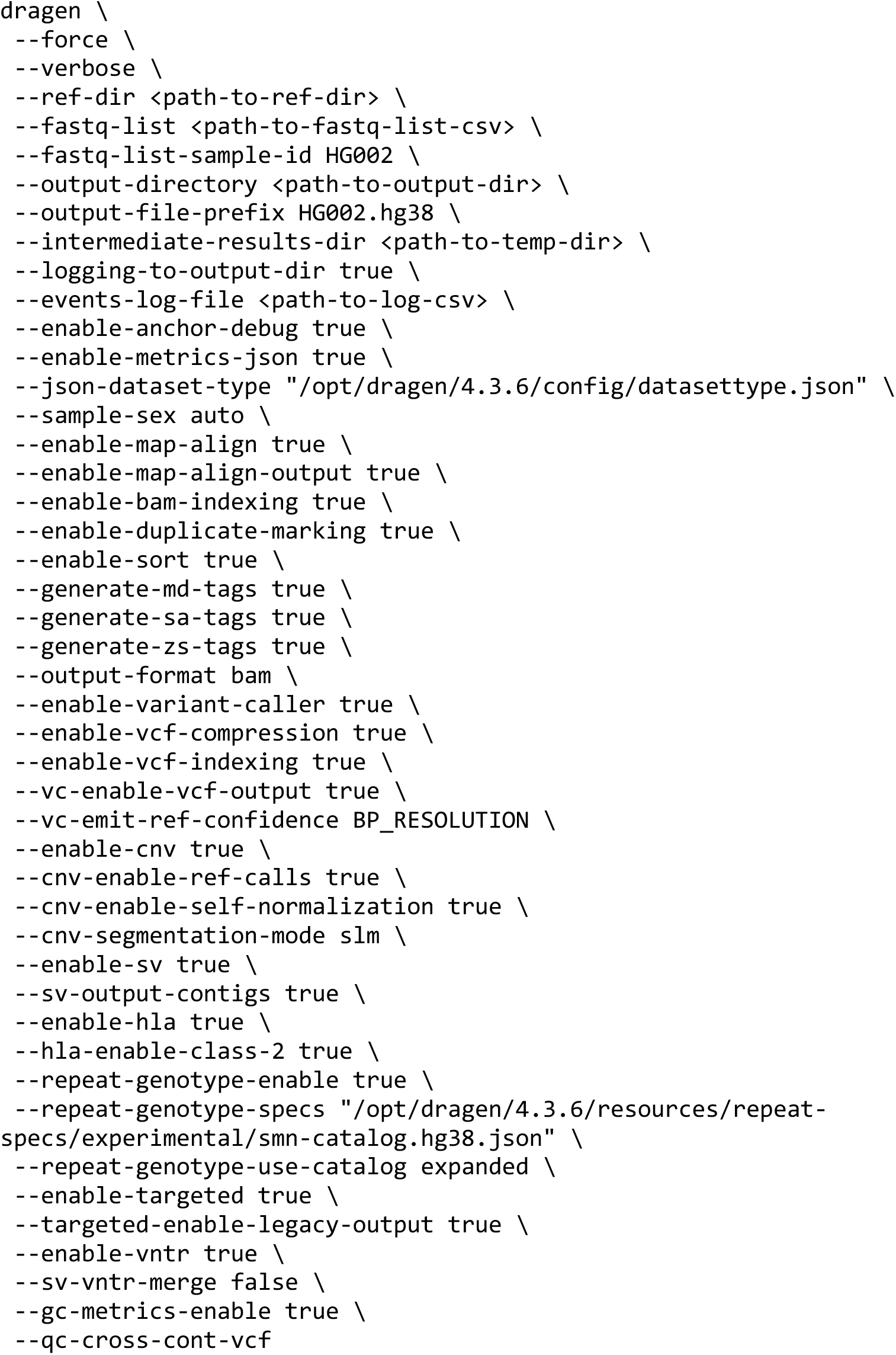
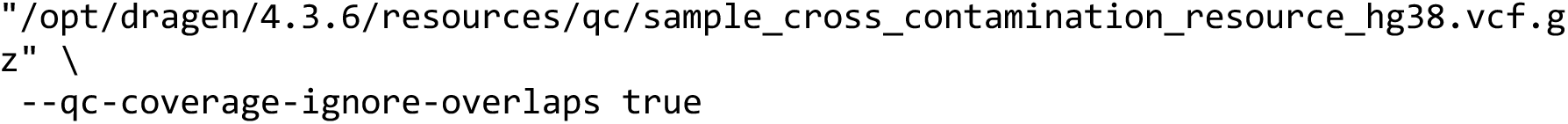

##### JHU ImputeFirst

Impute-first personalized pangenome pipeline, https://github.com/kvaddad1/impute-first. Input forward and reverse reads downloaded from https://storage.googleapis.com/brain-genomics-public/research/sequencing/fastq/novaseq/wgs_pcr_free/30x/HG002.novaseq.pcr-free.30x.R1.fastq.gz and https://storage.googleapis.com/brain-genomics-public/research/sequencing/fastq/novaseq/wgs_pcr_free/30x/HG002.novaseq.pcr-free.30x.R2.fastq.gz. The pipeline uses GRCh38 genetic maps in PLINK format (https://bochet.gcc.biostat.washington.edu/beagle/genetic_maps/plink.GRCh38.map.zip) and the HGSVC2 reference panel (https://ftp.1000genomes.ebi.ac.uk/vol1/ftp/data_collections/HGSVC2/release/v2.0/integrated_callset/). Read alignment and mapping are performed with `seqtk` (1.4-r122), `Bowtie2` (v2.5.4), `VG` (autoindex/giraffe v1.55.0 “Bernolda”), and SAMtools (1.21). The pipeline includes initial variant calling with `BCFtools` (v1.21), imputation with `Beagle` (v5.1 18May20.d20, Java), graph construction and alignment with `VG giraffe`, and variant calling with `DeepVariant` (v1.6.1). A summary of the pipeline as a bash script is available at https://gist.github.com/sinamajidian/c5a4aeb91e711930a48d3605d3c1d060.

NeuSomatic

The NeuSomatic variant callset was generated using the germline-adapted version of NeuSomatic and a model trained on multi-technology inputs (https://github.com/bioinform/neusomatic).^44^ Sequencing data from the PrecisionFDA truth challenge-v2 data^45^ were aligned to GRCh38 using BWA-MEM^46^ for Illumina, minimap2^47^ for PacBio, and ONT.

##### PacBio DeepVariant (Small Variants) and sawfish (SVs)

The HiFi-human-WGS-WDL v2.0.7 workflow (https://github.com/PacificBiosciences/HiFi-human-WGS-WDL/releases/tag/v2.0.7) was used for alignment and variant calling. HiFi PacBio reads generated SPRQ chemistry (https://downloads.pacbcloud.com/public/revio/2024Q4/WGS/GIAB_trio/HG002_rep1/, README).

Read were aligned using `pbmm2` v1.16.0 (https://github.com/PacificBiosciences/pbmm2/releases/tag/v1.16.0, container: quay.io/pacbio/pbmm2@sha256:24218cb5cbc68d1fd64db14a9dc38263d3d931c74aca872c998d12ef43 020ef0, task: https://github.com/PacificBiosciences/wdl-common/blob/60e9eb5c798b6d35dc26fe8561423cf1dab193bd/wdl/tasks/pbmm2.wdl). Small variants called with `DeepVariant`^6^ v1.6.1 (https://github.com/google/deepvariant/releases/tag/v1.6.1, container: https://hub.docker.com/layers/google/deepvariant/1.6.1/images/sha256-ccab95548e6c3ec28c75232987f31209ff1392027d67732435ce1ba3d0b55c68?context=explore, task: https://github.com/PacificBiosciences/wdl-common/blob/60e9eb5c798b6d35dc26fe8561423cf1dab193bd/wdl/workflows/deepvariant/deepvariant.wdl). SVs were called using `sawfish` v0.12.7 (https://github.com/PacificBiosciences/sawfish/releases/tag/v0.12.7)^19^ with the following commands.

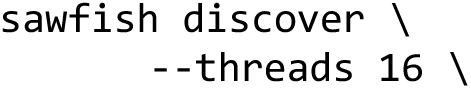
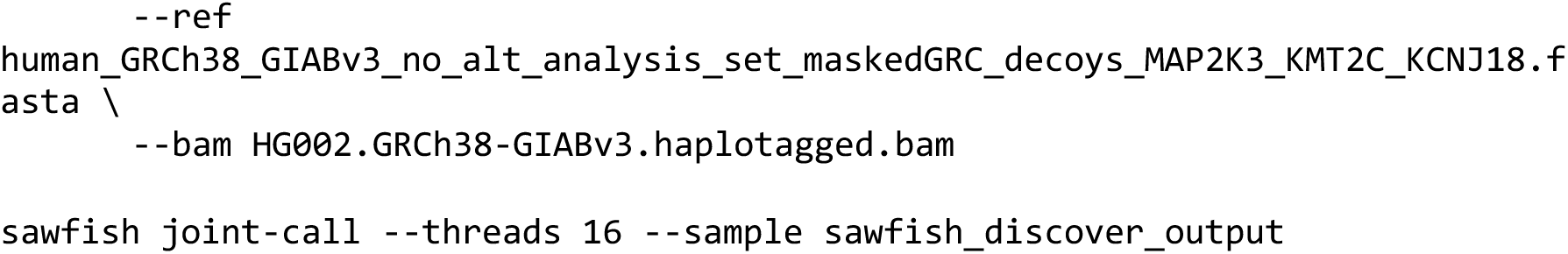

##### UCLA gSV

Germline SV calls for HG002 were made using `pipeline-call-gSV` v5.2.0 that uses `Delly` v1.2.6^54^ and `Manta` v1.6.0^55^ for SV calling (https://github.com/uclahs-cds/pipeline-call-gSV/tree/v5.2.0). Raw algorithm SV callsets were used to generate a consensus SV callset. Consensus SV calls were generated by comparing them to gnomAD-4.1.0 SV data using `BCFtools-1.15.1 isec` with `-- nfiles +1`. Additionally, SVs uniquely called by `Delly` v1.2.6 and `Manta` v1.6.0 were filtered to retain only PASS calls.

##### Ultima Genomics

Variant callset generated using Ultima’s aligner and an optimized version of DeepVariant. Input data^48^ available at s3://ultimagen-feb-2024-giab/Crams/NA24385-Z0027.cram.

Alignment was performed using Ultima’s aligner, `UA`, v5.1.0.19 (10b91e1). Reference genome: gs://gcp-public-data–broad-references/hg38/v0/Homo_sapiens_assembly38.fasta. Ultima’s alignment tool on AWS HealthOmics: https://github.com/Ultimagen/healthomics-workflows/blob/main/workflows/trim_align_sort/trim_align_sort.md. Variant calling was performed using UG’s optimized somatic `DeepVariant` v1.4. Ultima’s somatic caller on AWS HealthOmics: https://github.com/Ultimagen/healthomics-workflows/blob/main/workflows/efficient_dv/howto-germline-calling-efficient-dv.md. See the AppNote for additional details (https://cdn.sanity.io/files/l7780ks7/production/d7e5bf926016a02f682f10415e45224f75e307a3.pdf.)

#### External Evaluation Manual Curation and Confidence Intervals

The GRCh38 version of the draft small and SV benchmark sets was compared small and SV callsets from external groups to ensure the benchmark set adheres to the RIDE principles (Fig. 1B and C). `hap.py,` with `vcfeval` as the comparison engine, was used to identify variant-call discrepancies, and `truvari` was used for SVs. Discrepancies were grouped into strata by variant type and genomic context, and a random subset from each stratum was selected for manual curation by the group that submitted the variant callset.

Discrepancies between the benchmark and query callsets were categorized into 14 strata for small variants and 8 for SVs. Five variants were randomly selected from each stratum for manual curation. Small-variant strata were defined by whether they were unique to the benchmark or query callset (discrepancy type), variant type (SNP or INDEL), and genomic context. Within each discrepancy-type and variant-type branch, strata were assigned sequentially to ensure mutually exclusive classification. SNPs were prioritized by tandem-repeat overlap, difficult-to-map/repetitive context, XY non-pseudoautosomal location, and residual context. INDELs were prioritized by homopolymers longer than 6 bp, tandem-repeat overlap, difficult-to-map/repetitive context, XY non-pseudoautosomal location, and residual context. Difficult-to-map/repetitive context included low-mappability annotations, segmental duplications, and LINE, LTR, or satellite `RepeatMasker` annotations. Five variants per stratum were randomly sampled using a fixed random seed. XY non-pseudoautosomal strata were excluded from the final exported manual-curation set. SV discrepancies were similarly categorized into 8 strata based on discrepancy type, variant type (insertion or deletion), and whether the variant was within a tandem repeat.

Using the `Miqa` bioinformatics platform (Magna Labs), evaluators classified the discrepancies as correct, incorrect, or unsure based on alignments for multiple sequencing technologies (Illumina 2 × 250 bp, Element short and long insert data, PacBio HiFi, PacBio Onso, and ONT ultra-long) for the draft benchmark set and their callsets. For each variant, evaluators were presented with a pre-configured multi-track, multi-genome `Miqa` session displaying alignments via the Integrative Genomics Viewer (IGV) to GRCh38 and the HG002 Q100 v1.1 maternal and paternal haplotype assemblies, with navigation and display context tailored to each variant’s type and size across each reference. Curations were recorded independently for each external group. Variants identified as incorrect or unsure in the draft benchmark set were re-curated by the NIST team, and curations confirmed by the group that submitted the callsets. Small variant evaluation subset table generation pipeline https://github.com/nate-d-olson/giab-smvar-curation-table and SV evaluation subset table generation pipeline https://github.com/nate-d-olson/giab-stvar-curation-table.

To estimate the probability that a discrepancy between the benchmark set and a comparison callset represents an error in the comparison callset (i.e., the benchmark is correct), we fit separate binomial generalized linear models (GLMs) with a logit link function for small variants and structural variants. Manually curated discrepancies were classified as “correct in benchmark” (yes), “not correct in benchmark” (no), or “unsure”. The primary analysis treated variants curated as “unsure” and “no” as failures (i.e., the discrepancy is an error in the benchmark), using this as a lower bound for the probability estimate. For an upper bound, variants curated as “unsure” and “yes” were treated as successes (the discrepancy is due to an error in the comparison callset).

For small variants (N = 350; 5 callsets), the model included three additive predictors: variant type (SNP vs. INDEL), genomic context (autosome, tandem repeat, homopolymer > 6 bp, or low-mappability region), and whether the discrepancy was unique to the query callset or benchmark set. For SVs (N = 280; 7 callsets), predictors were repeat annotation (TR vs. non-TR), SV category (DEL vs. INS), and unique-to status (comparison vs. benchmark). All predictors were entered as main effects without interactions. As different sets of variants were curated for each callset, each observation represents an independent evaluation at a distinct genomic position, and the callset was not included as a model term.

To establish a lower bound on how reliably the benchmark identifies errors in comparison callsets, one-sided 95 % lower confidence bounds were reported rather than two-sided intervals.

Marginal probabilities and confidence intervals on the probability scale were obtained using the `emmeans` R package (2.0.4) with proportional weighting to average over the other model terms. Proportional weighting ensures that factor levels are weighted by their observed frequencies in the data, which is relevant because the homopolymer > 6 bp context applies only to INDELs. In the sensitivity analysis, the small variant model was fit using bias-reduced logistic regression (Firth penalization via the `brglm2` R package, version 1.1.0) because treating “unsure” as correct left only 5 failures across 350 observations, resulting in quasi-complete separation in the low mappability stratum.

