## Supplementary figures and images for "The Genome In A Bottle HG002 assembly-based variant benchmark set enables comprehensive benchmarking of small and structural variants"

### Supplemental Figure

#### HG002 Q100 Variant Benchmark − GRCh38 Region Coverage

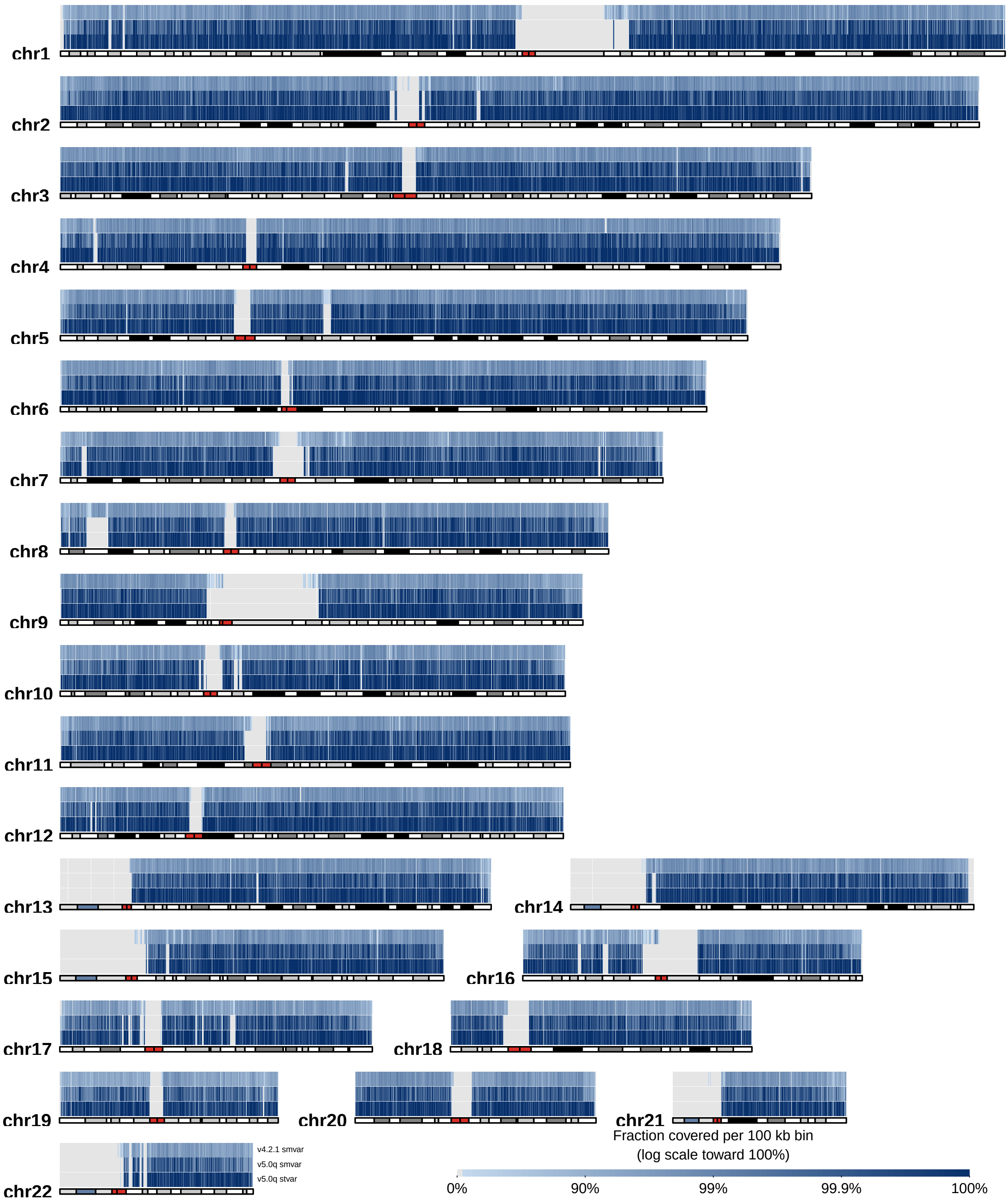
